# Reconstituted depolarization-induced Ca^2+^ release platform for skeletal muscle disease mutation validation and drug discovery

**DOI:** 10.1101/2022.06.29.498090

**Authors:** Takashi Murayama, Nagomi Kurebayashi, Takuro Numaga-Tomita, Takuya Kobayashi, Satoru Okazaki, Kyosuke Yamashiro, Tsutomu Nakada, Shuichi Mori, Ryosuke Ishida, Hiroyuki Kagechika, Mitsuhiko Yamada, Takashi Sakurai

## Abstract

In skeletal muscle excitation-contraction (E-C) coupling, depolarization of the plasma membrane triggers Ca^2+^ release from the sarcoplasmic reticulum (SR), referred to as depolarization-induced Ca^2+^ release (DICR). DICR occurs via the type 1 ryanodine receptor (RyR1), which physically interacts with the dihydropyridine receptor Cav1.1 subunit in specific machinery formed with additional essential components including β1a, Stac3 adaptor protein and junctophilins. Exome sequencing has accelerated the discovery of many novel mutations in genes encoding DICR machinery in various skeletal muscle diseases. However, functional validation is time-consuming because it must be performed in a skeletal muscle environment. In this study, we established a platform of the reconstituted DICR in HEK293 cells. The essential components were effectively transduced into HEK293 cells expressing RyR1 using baculovirus vectors, and Ca^2+^ release was quantitatively measured with R-CEPIA1er, a fluorescent ER Ca^2+^ indicator, without contaminant of extracellular Ca^2+^ influx. In these cells, [K^+^]-dependent Ca^2+^ release was triggered by chemical depolarization with the aid of inward rectifying potassium channel, indicating a successful reconstitution of DICR. Using the platform, we evaluated several Cav1.1 mutations that are implicated in malignant hyperthermia and myopathy. We also tested several RyR1 inhibitors; whereas dantrolene and Cpd1 inhibited DICR, procaine had no effect. Furthermore, twitch potentiators such as perchlorate and thiocyanate shifted the voltage dependence of DICR to more negative potentials without affecting Ca^2+^-induced Ca^2+^ release. These results well reproduced the findings with the muscle fibers and the cultured myotubes. Since the procedure is simple and reproducible, the reconstituted DICR platform will be highly useful for validation of mutations and drug discovery for skeletal muscle diseases.

**Summary:** Mutations in essential components for depolarization-induced Ca^2+^ release (DICR) are implicated into various skeletal muscle diseases. Murayama et al. establish a reconstituted DICR platform in nonmuscle cells for evaluation of disease-causing mutations and drug discovery.

## Introduction

In vertebrate skeletal muscle, depolarization of the plasma membrane triggers Ca^2+^ release from the sarcoplasmic reticulum (SR), which is commonly referred to as excitation-contraction (E-C) coupling. This process occurs at triad junctions in which a transverse (T) tubule is closely apposed to terminal cisterna of the SR (Franzini-Armstrong, 2018). In excitation-contraction coupling, the type 1 ryanodine receptor (RyR1), a Ca^2+^ release channel in the SR, is activated by a conformational change of the voltage sensor in the Cav1.1 subunit of the dihydropyridine receptor (DHPR) upon depolarization of the T-tubule membrane. This is referred to as depolarization-induced Ca^2+^ release (DICR) (Rios and Pizarro, 1991; Schneider, 1994). Studies with knockout animals have revealed that functional and structural interactions between Cav1.1 and RyR1 in muscle require at least three additional components, a β1a auxiliary subunit of DHPR (Gregg et al., 1996; Schredelseker et al., 2005), the Stac3 adaptor protein (Horstick et al., 2013; Nelson et al., 2013; Reinholt et al., 2013), and junctophilins (Takeshima et al., 2000; Yang et al., 2022), which may form specific machinery for DICR (Flucher and Campiglio, 2019; Shishmarev, 2020; Rufenach and Van Petegem, 2021).

Mutations in genes that encode DICR machinery components have been implicated in various types of muscle diseases, including congenital myopathies, periodic paralysis, and malignant hyperthermia (MH) (Agrawal et al., 2018; Flucher, 2020; Lawal et al., 2020). Rapid exome sequencing has accelerated the discovery of novel mutations; however, validation of these mutations by functional testing is critically important for the diagnosis and treatment of these diseases. This has been performed using myotubes from mice that lack DICR machinery components, for example knockout mice of Cav1.1 (Tanabe et al., 1988), β1a (Gregg et al., 1996) and Stac3 (Nelson et al., 2013). While this approach has evaluated several disease-causing mutations in the DICR components (Weiss et al., 2004; Pirone et al., 2010; Eltit et al., 2012; Polster et al., 2016; Perez et al., 2018), it has certain limitations. For example, it is expensive and labor intensive to maintain the mouse lines. Differentiation of cells into myotubes is time consuming and is affected by various factors, such as the cell line used and experimental conditions. Furthermore, transduction of foreign genes into myotubes is often inefficient.

There are no specific treatments for most diseases related to mutations in the DICR machinery. A major cause for the slow pace of novel drug development is the lack of an appropriate drug screening platform for these diseases. We have recently established an efficient high-throughput platform for screening RyR1-targeted drugs using endoplasmic reticulum (ER) Ca^2+^ measurements in HEK293 cells (Murayama et al., 2018; Murayama and Kurebayashi, 2019) and using this platform we developed Compound 1 (Cpd1), a novel RyR1 inhibitor for treatment of MH (Mori et al., 2019; Yamazawa et al., 2021). Therefore, functional screening using HEK293 cells is a promising platform for drug discovery.

Recently, Perni et al. successfully reconstituted DICR by co-expressing five essential components, RyR1, Cav1.1, β1a, junctophilin-2 (JP2) and Stac3, in non-muscle tsA201 cells (Perni et al., 2017). They demonstrated formation of ER-plasma membrane junctions in which Cav1.1 is arranged into tetrads, which is indicative of physical links to RyR1 (Block et al., 1988). Depolarization of the plasma membrane using the patch-clamp technique induced a rapid Ca^2+^ transient without influx of extracellular Ca^2+^. This provides a basis for developing reconstituted DICR.

In this study, we established a platform for DICR that was reconstituted in HEK293 cells. Baculovirus (BV) infection of essential components greatly improved transduction efficiency. Chemical depolarization by high concentration potassium [K^+^] solutions with the aid of an inward-rectified potassium channel (Kir2.1) enabled simultaneous stimulation of many cells. Furthermore, a genetically-encoded ER Ca^2+^ indicator quantitatively measured DICR without influence of extracellular Ca^2+^ influx. Using this platform, we successfully evaluated disease-causing mutations in Cav1.1 and tested several RyR1 inhibitors and DICR modulators. Our reconstituted DICR platform will accelerate the validation of mutations and drug discovery for skeletal muscle diseases.

## Materials and Methods

### Generation of stable inducible HEK293 cell lines

HEK293 cells stably expressing R-CEPIA1er and RyR1 were generated as described previously (Murayama et al., 2018; Murayama and Kurebayashi, 2019). Briefly, a full-length rabbit RyR1 cDNA was cloned in a tetracycline-induced expression vector (pcDNA5/FRT/TO; Life Technologies, CA, USA) (Tong et al., 1997). Flp-In T-REx HEK293 cells were co-transfected with this expression vector and pOG44 in accordance with the manufacturer’s instructions. Clones with suitable doxycycline-induced expression of RyR1 were selected and used for experiments. The cells were then transfected with an R-CEPIA1er cDNA (a generous gift from Masamitsu Iino, The University of Tokyo) for stable expression of R-CEPIA1er. Cells were cultured in Dulbecco’s modified Eagle’s medium supplemented with 10% fetal calf serum, 2 mM L-glutamine, 15 µg/ml blasticidin, and 100 µg/ml hygromycin.

### BV Production

Baculovirus (BV) that expresses vesicular stomatitis virus G-protein (VSVG) on the virus envelope (Tani et al., 2001) was produced using the Bac-to-Bac system (LifeTechnologies) as described previously (Uehara et al., 2017). cDNAs for Cav1.1, β1a, JP2, Stac3, and Kir2.1 were cloned from mouse skeletal muscle using reverse transcription-PCR (primer pairs are listed in **Table 1**). Ca^2+^ non-conducting mutation (N617D) (Schredelseker et al., 2010) and disease-causing mutations were introduced in Cav1.1 cDNA by inverse PCR (primer pairs are listed in **Table 1**). Each cDNA was cloned into the modified donor vector (pFastBac1-VSVG-CMV-WPRE). BV was produced in Sf9 cells according to the manufacturer’s instructions (LifeTechnologies). BV titers were immunologically determined using an anti-gp64 antibody (AcV5, Santa Cruz Biotechnology)(Kitts and Green, 1999). Titers were 1–2×10^8^ pfu/mL for all viruses. The BV solutions were stored in the dark at 4°C and used within one month after production.

**Table 1.**
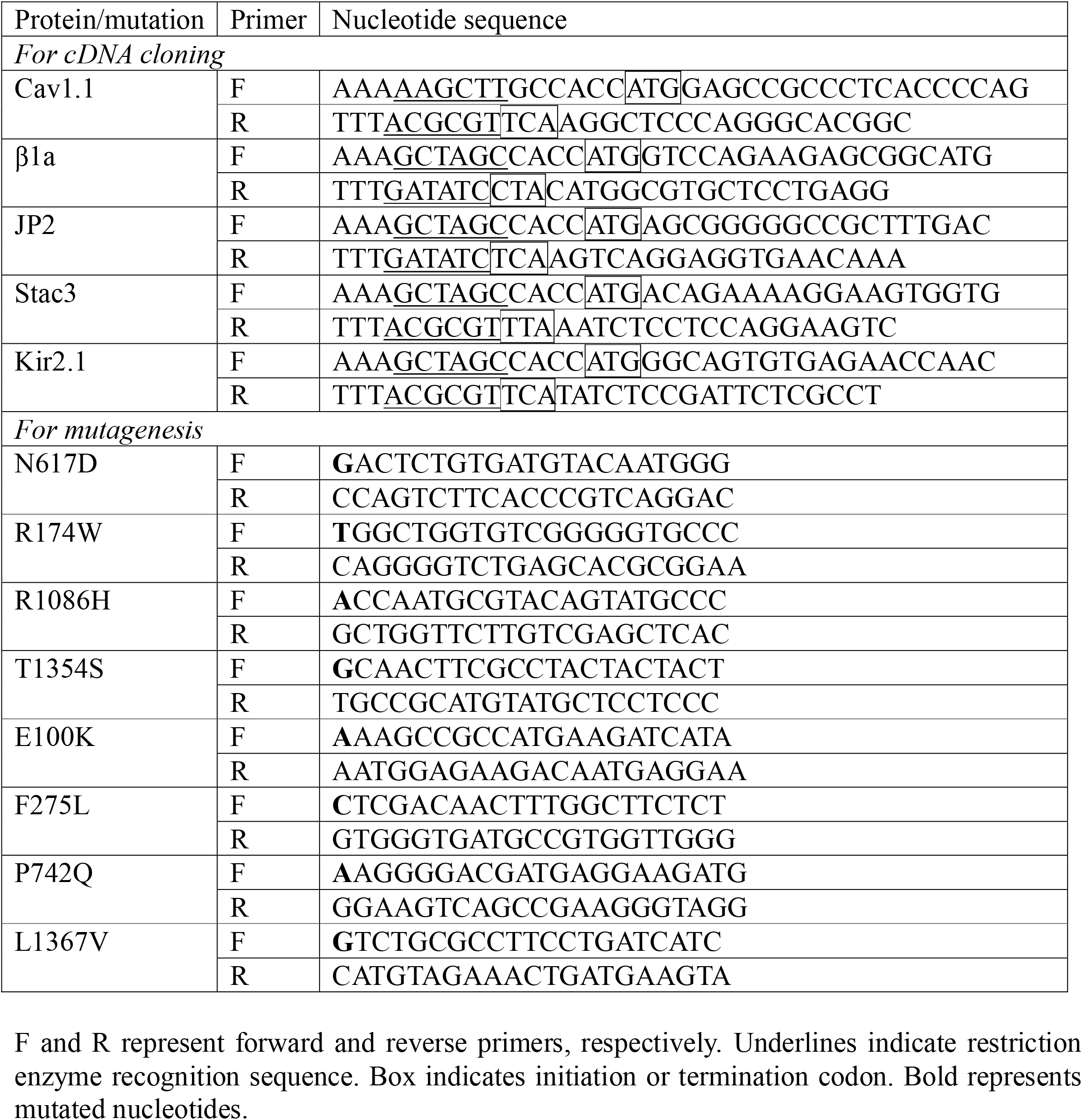
List of PCR primers.

### Immunocytochemistry

RyR1-HEK293 cells were seeded on a 35 mm glass bottom dish that had been coated with collagen solution at a density of 2×10^5^ cells/well in 1.5 mL culture media. One day after seeding, culture media were exchanged to those containing 2 µg/mL doxycycline (for induction of RyR1) and BV solutions for Cav1.1 (N617D), β1a, JP2, Stac3, and Kir2.1 (30 µL each in 1.5 mL). Multiplicity of infection (MOI) was 7.5-15 for each BV (depending on the virus titer). After 24 h, cells were fixed with ice-cold methanol and rinsed with phosphate-buffered saline (PBS). The cells were blocked with 5% BSA/PBS for 30 min and incubated overnight with primary antibodies against Cav1.1 (sc-514685, Santa Cruz Biotechnology), β1a (MA5-27714, Invitrogen), JP2 (sc-377086, Santa Cruz Biotechnology), Stac3 (1G8, Abnova), or Kir2.1 (S112B-14, Invitrogen) in 1% BSA/PBS. After washing three times with PBS, the cells were incubated for 2 h with Alexa594-labeled goat anti-mouse IgG (Invitrogen) in 1% BSA/PBS. After washing with PBS, the cells were observed using an all-in-one fluorescence microscope (BZ-X710, Keyence, Osaka, Japan).

### Single-cell Ca^2+^ imaging

RyR1/R-CEPPIA1er HEK293 cells were seeded on a 35 mm glass bottom dish and culture media were exchanged next day as described above. 24 h after infection, single-cell Ca^2+^ imaging was performed as described previously (Murayama et al., 2015; Uehara et al., 2017). Solutions were perfused using an in-line solution heater/cooler (Warner Instruments, Holliston, MA, USA). Measurements were performed at 26°C instead of 37°C, since cells tended to detach from the dish during perfusion at 37°C. ER [Ca^2+^] signals were determined by R-CEPIA1er fluorescence excited at 561 nm using a 20× objective lens. Cells were perfused with normal Krebs solution (140 mM NaCl, 5 mM KCl, 2 mM CaCl_2_, 1 mM MgCl_2_, 11 mM glucose, 10 mM HEPES, pH 7.4) for 30 s, high [K^+^] Krebs solution containing 50 mM [K^+^] for 50 s, and normal Krebs solution for 1.5 min. Fluorescence intensities in individual cells were determined by region of interest (ROI) analysis using AquaCosmos software (Hamamatsu Photonics, Hamamatsu, Japan).

### Western blotting

RyR1-HEK293 cells were seeded on 12-well plate at a density of 3×10^5^ cells/well in 1 mL culture media. One day after seeding, 1 mL culture media containing 2 µg/mL doxycycline (for induction of RyR1) and BV solutions for Cav1.1 (N617D), β1a, JP2, Stac3, and Kir2.1 (20 µL each) were added to each well. After 24 h, cells were harvested, rinsed with PBS and lysed with 50 µL of Pro-Prep protein extraction solution (iNtRON Biotechnology). After centrifugation at 15,000 rpm for 5 min, the extracted proteins were separated on 3–15% polyacrylamide gels and transferred to PVDF membranes. Membranes were probed with the primary antibodies described above and calnexin (C4731, Sigma-Aldrich, MO, USA) as a loading control, followed by HRP-labeled anti-mouse IgG (04-18-18, KPL) or anti-rabbit IgG (074-1516, KPL). Positive bands were detected by chemiluminescence using ImmunoStar LD (Fujifilm Wako Chemicals) as a substrate.

### Time lapse ER [Ca^2+^] Measurements using a fluorometer

Time lapse ER [Ca^2+^] measurements were performed using the FlexStation3 fluorometer (Molecular Devices, San Jose, CA) (Murayama et al., 2018; Murayama and Kurebayashi, 2019). Stable RyR1/R-CEPIA1er HEK293 cells were seeded on 96-well flat-, clear-bottom black microplates (#3603, Corning, New York, NY) at a density of 3×10^4^ cells/well in 100 µL culture media. One day after seeding, 100 µL culture media containing 2 µg/mL doxycycline (for induction of RyR1) and BV solutions for Cav1.1 (N617D), β1a, JP2, Stac3, and Kir2.1 (2 µL each) were added to each well. Multiplicity of infection (MOI) was 7.5-15 for each BV (depending on the virus titer). After 24 h, the culture media were replaced with 81 µL normal Krebs solution, and the microplate was placed in a FlexStation3 fluorometer preincubated at 37°C. Signals from R-CEPIA1er, which was excited at 560 nm and emitted at 610 nm, were captured every 5 sec for 150 sec. Thirty sec after starting, 54 µL of the high [K^+^] or caffeine solution was applied to the cells. The fluorescence change was expressed as F/F_0_ in which averaged fluorescence intensity of the last 25 sec (F) was normalized to that of the initial 25 sec (F_0_). For drug testing, compounds were added to normal Krebs solution at the indicated concentrations.

For determination of resting ER [Ca^2+^], cells were incubated with 81 µL normal Krebs solution, and 54 µL Krebs solution containing 50 mM Ca^2+^ and 50 µM ionomycin was added 30 sec after starting (final concentrations of Ca^2+^ and ionomycin were 20 mM and 20 µM, respectively). R-CEPIA1er fluorescence was captured every 10 sec for 6 min.

### Measurement of resting membrane potential

The resting membrane potential of BV-infected RyR1-HEK293 cells with or without Kir2.1 expression was measured under current-clamp conditions in the whole cell configuration of the patch-clamp method with an Axopatch 200B amplifier (Molecular Devices, San Jose, CA). Pipettes were fabricated from borosilicate glass capillaries (Kimble) with a vertical pipette puller (PP-83, Narishige, Tokyo, Japan). Pipettes had series resistance of 3–4 MΩ when filled with the pipette solution containing (in mM) 120 D-glutamate, 20 KCl, 10 NaCl, 0.1 EGTA, 2 MgCl_2_, 3 MgATP, and 5 HEPES (pH 7.4 adjusted with KOH). The series resistance was routinely compensated by ∼75% with the amplifier. Bath solution contained (in mM): 140 NaCl, 5 KCl, 11 glucose, 2 CaCl_2_, 1 MgCl_2_, and 10 HEPES (pH 7.4 adjusted with NaOH). High [K^+^] bath solution contained (in mM): 95 NaCl, 50 KCl, 11 glucose, 2 CaCl_2_, 1 MgCl_2_, and 10 HEPES (pH 7.4 adjusted with NaOH). The resting membrane potential was measured immediately after breaking into cells.

### Statistics

Data are presented as means ± SD. Unpaired Student’s t test was used for comparisons between two groups. One-way analysis of variance (ANOVA), followed by Dunnett’s test, was performed to compare three or more groups. Two-tailed tests were used for all analyses. Statistical analysis was performed using Prism v9 (GraphPad Software, Inc., La Jolla, USA).

## Results

### Establishment of a reconstituted DICR platform using a fluorescence microplate reader

To reconstitute DICR, we used HEK293 cells stably expressing RyR1 and R-CEPIA1er, a genetically-encoded fluorescent ER Ca^2+^ indicator (Suzuki et al., 2014), that were generated for high-throughput screening of novel RyR1 inhibitors (Murayama et al., 2018; Murayama and Kurebayashi, 2019) (**Fig. 1 A**, left). The other components for DICR, Cav1.1, β1a, Stac3 and JP2, which were used by Perni et al. (Perni et al., 2017), were transduced using VSV-G pseudotyped baculovirus (BV), which efficiently infects mammalian cells (Tani et al., 2001; Uehara et al., 2017) (**Fig. 1 A**, right). In these cells, DICR can be detected by a decrease in ER Ca^2+^. To prevent extracellular Ca^2+^ influx through Cav1.1 by depolarization, a N617D mutation was introduced that abolishes Ca^2+^ conductance (Schredelseker et al., 2010). In addition, we expressed Kir2.1, an inward-rectified potassium channel in skeletal muscle (DiFranco et al., 2015), to hyperpolarize the membrane potential because the resting membrane potential in HEK293 cells is much less (^−^20 mV) than in skeletal muscle (−90 mV) (Kirkton and Bursac, 2011). Imunocytochemistry revealed that the five proteins (Cav1.1, β1a, Stac3, JP2, and Kir2.1) were successfully expressed in almost all cells with infection of 5 BV, which we term hereafter 5 BV cells, but not in non-infected cells (No BV cells) (**Fig. S1 A**). BV-dependent protein expression was also confirmed by Western blotting of cell lysate (**Fig. S1 B**). Therefore, BV infection effectively transduced the essential components of DICR in RyR1/R-CEPIA1er-HEK293 cells.

**Figure 1.**
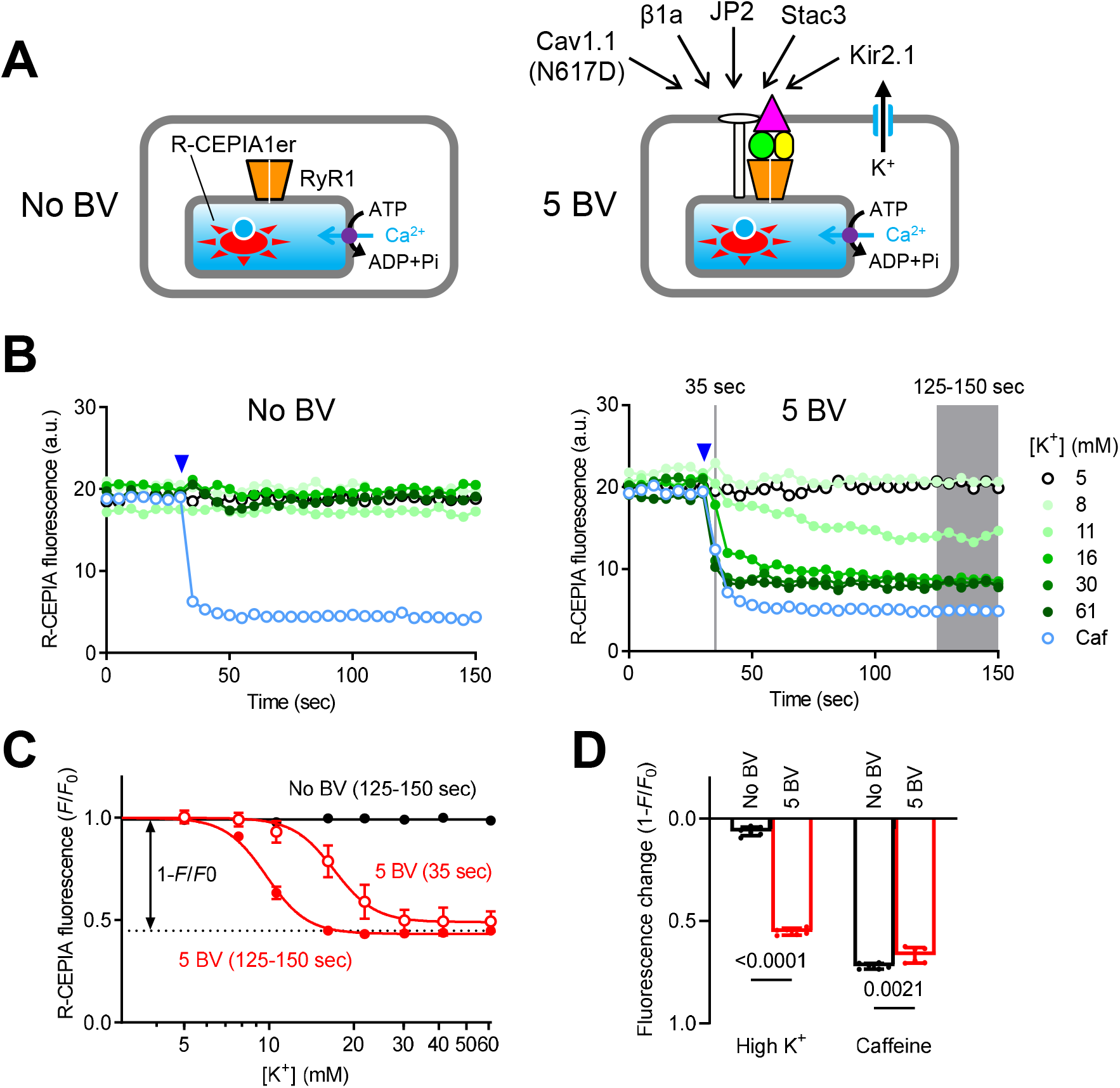
DICR measurements in 96-well plates using a fluorescence microplate reader. **(A)** Schematic drawing of RyR1/R-CEPIA1er HEK293 cells without (No BV, left) and with (5 BV, right) BV infection of essential components for DICR (Cav1.1 carrying N617D Ca^2+^ non-conducting mutation, β1a, JP2, Stac3, and Kir2.1). **(B)** Typical results of time-lapse R-CEPIA1er fluorescence measurement using a FlexStation3 fluorescence microplate reader for RyR1/R-CEPIA1er HEK293 cells without (left, No BV) or with (right, 5 BV) BV infection. High [K^+^] solution ranging from 5–61 mM (symbols shown in right) or 10 mM caffeine (light blue) was applied at 30 sec after starting (blue arrowheads). **(C)** [K^+^] dependence of R-CEPIA1er fluorescence (F/F_0_) in No BV (black) and 5 BV cells (red). F/F_0_ was obtained by normalizing the fluorescence (F) at 35 sec (open circles) or the averaged fluorescence of 125-150 sec to that of the first 25 sec (F_0_). Data are means ± SD (n = 9, *N* = 3 for 5 BV and n = 6, *N* = 3 for No BV). Note that [K^+^] dependent plot with data at 35 sec (open red) was shifted rightward compared with that from data at 125–150 sec (filled red) with larger variations, especially around EC_50_ values. **(D)** Fluorescence change by 61 mM [K^+^] or 10 mM caffeine. Fluorescence change is expressed as 1-F/F_0_ (see panel **C**). Data are means ± SD (n = 6, *N* = 3) and were analyzed by two-way ANOVA with Bonferroni’s multiple comparisons test. Note that 5 BV cells show large fluorescence change with high [K^+^] which is nearly 80% of that with caffeine. “n” is the number of wells and “*N*” is the number of independent experiments.

To detect DICR, we used a fluorescence microplate reader (FlexStation3), which measures time-lapse R-CEPIA1er fluorescence in 96-well plates. Fluorescence signals were collected in a 1.5 mm diameter area; therefore, an averaged signal from thousands of cells was detected in each well. We adopted chemical depolarization using high [K^+^] solutions, which can simultaneously depolarize all the cells to trigger DICR. The cells were initially incubated with normal Krebs solution containing 5 mM [K^+^]. Then, solutions containing various [K^+^] concentrations were added to give [K^+^] concentrations of 5–61 mM. In cells without BV infection, no apparent changes in fluorescence were observed with any [K^+^] concentration (**Fig. 1 B**, left). Caffeine (10 mM) reduced the R-CEPIA1er fluorescence, confirming the functional expression of RyR1. In 5 BV cells, high [K^+^] solution rapidly reduced the R-CEPIA1er fluorescence in a [K^+^]-dependent manner (**Fig. 1 B**, right). To monitor ER Ca^2+^ changes in individual cells, we also observed R-CEPIA1er fluorescence using laser scanning confocal microscopy. In 5 BV cells, high [K^+^] solution caused a large and transient reduction in the R-CEPIA1er fluorescence in >90% of cells (**Movie S1** and **Fig. S2**).

For quantitative analysis, we normalized the averaged fluorescence intensity to that of the initial 25 sec (F_0_) to correct variations in the initial fluorescence intensity between wells. We first used rapid fluorescence change 5 sec after stimulation (i.e., 35 sec), which may reflect the rate of Ca^2+^ release. The F/F_0_ plot as a function of [K^+^] concentrations demonstrated [K^+^] dependence with an EC_50_ of 17.0 ± 0.4 mM. However, variations were relatively large, especially near EC_50_ value because of rapid changes in fluorescence intensity (**Fig. 1 C**). We instead adopted the average fluorescence intensity for the last 25 sec (125-150 sec). This gave an EC_50_ of 9.8 ± 0.1 mM [K^+^], which was lower than that obtained from 35 sec, but with smaller variations (**Fig. 1 C**). The fluorescence change by 61 mM [K^+^], which is expressed as 1-F/F_0_ (**Fig. 1 C**), reached ∼80% of that induced by caffeine in 5 BV cells, indicating a high degree of DICR reconstitution (**Fig. 1 D**). Taken together, these results indicate successful establishment of a reconstituted DICR platform.

The original method for DICR reconstitution monitors cytoplasmic [Ca^2+^] (Perni et al., 2017); therefore, we also assessed DICR by measuring cytoplasmic [Ca^2+^] using G-GECO1.1, a genetically-encoded Ca^2+^ indicator (Zhao et al., 2011). In cells without BV infection, a substantial increase in [Ca^2+^] was observed by high [K^+^], indicating the existence of endogenous Ca^2+^ influx pathways by depolarization (**Fig. S3 A**). In 5 BV cells, DICR was detected by rapid and large increases in [Ca^2+^], which declined with time (**Fig. S3 B**). The rapid fluorescence change 5 sec after stimuli (i.e., 35 sec) provided dose-dependent curve with EC_50_ of 16.4 ± 0.5 mM, which is similar to the value determined by rapid fluorescence change in ER [Ca^2+^] measurement (17 mM, see **Fig. 1 C**) and those obtained from mouse skeletal muscle myotubes (∼21 mM) in high [K^+^]-induced Ca^2+^ transients (Yang et al., 2006; Lopez et al., 2018).

### Properties of the reconstituted DICR

We next examined properties of the reconstituted DICR in HEK293 cells. Initially, we examined the role of the essential components. Removal of Cav1.1 resulted in complete loss of high [K^+^]-induced Ca^2+^ release (**Fig. 2 A-C**). Ca^2+^ release was also lost in cells lacking β1a or Stac3, and in cells expressing RyR2 instead of RyR1 (**Fig. 2 C**). These results are consistent with previous findings of mice lacking each component that they are essential for DICR (Tanabe et al., 1988; Takeshima et al., 1994; Gregg et al., 1996; Nelson et al., 2013). Interestingly, DICR still occurred without JP2, although the fluorescence change by 61 mM [K^+^] was significantly reduced and [K^+^] dependence was largely shifted rightward (**Fig. 2 B, C**).

**Figure 2.**
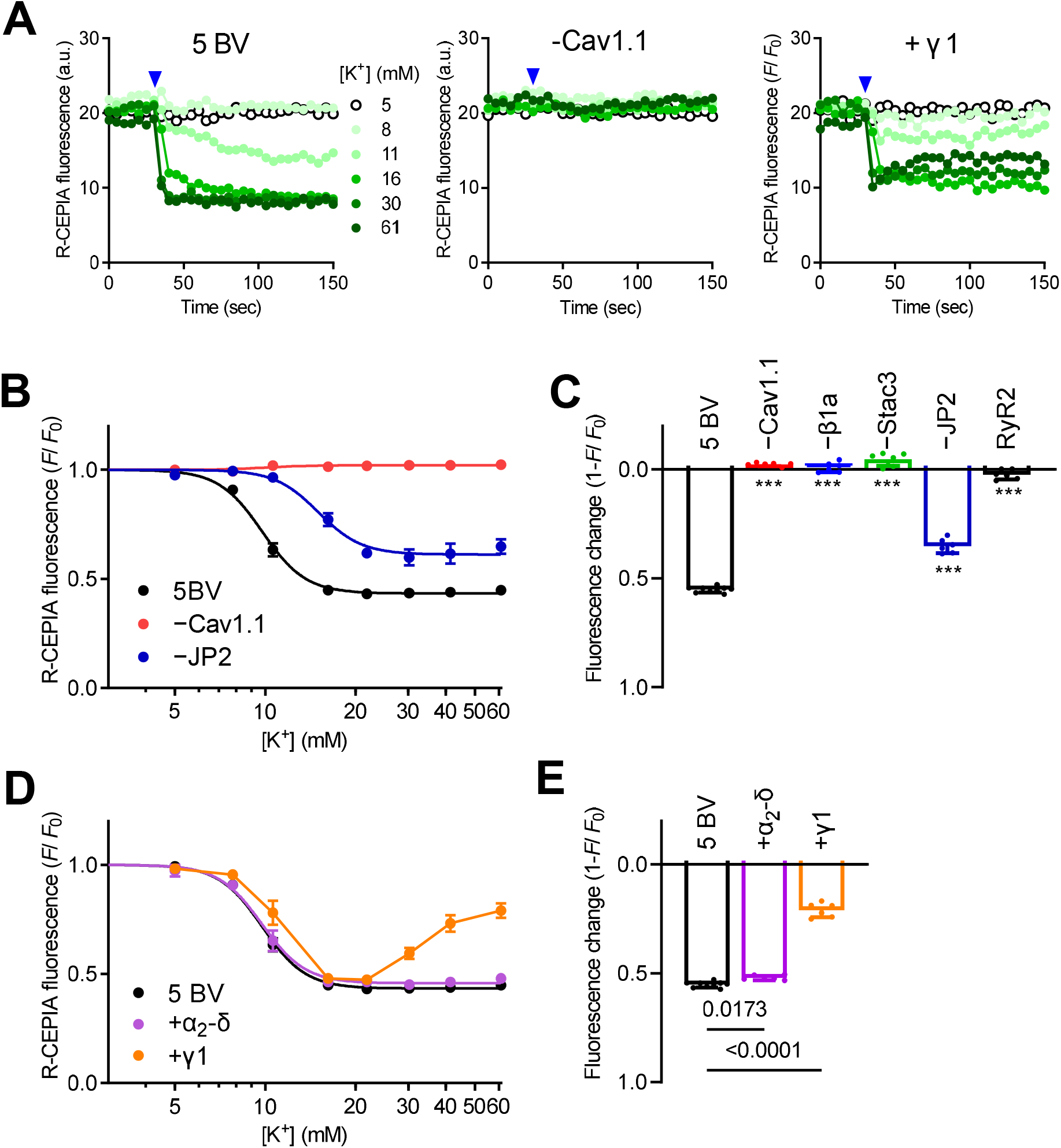
Properties of the reconstituted DICR platform. **(A)** Typical results of R-CEPIA1er fluorescence in cells infected with 5 BV (left), BV without Cav1.1 (center), and 5 BV plus γ1 (right). (**B**) [K^+^] dependence of R-CEPIA1er fluorescence (F/F_0_) in 5 BV (black), -Cav1.1 (red), and -JP2 (dark blue). Data are means ± SD (n = 9, *N* = 3 for 5 BV; n = 6, *N* = 3 for -Cav1.1 and -JP2). **(C)** Fluorescence change by 61 mM [K^+^] in the BV-infected cells lacking one of essential components (−Cav1.1, -β1a, -Stac3, or -JP2) or with RyR2 cells instead of RyR1. Data are means ± SD (n = 9, *N* = 3 for 5 BV and n = 6, *N* = 3 for the others) and were analyzed by one-way ANOVA with Dunnett’s multiple comparisons test. ^***^, p < 0.0001 from 5 BV. **(D)** [K^+^] dependence of R-CEPIA1er fluorescence (F/F_0_) in cells infected with 5 BV (black), 5 BV plus α_2_-δ (purple) and 5 BV plus γ1 (orange). Data are means ± SD (n = 9, *N* = 3 for 5 BV; n = 6, *N* = 3 for + α_2_-δ and + γ1). **(E)** Fluorescence change by 61 mM [K^+^] in 5 BV, +α_2_-δ, and +γ1 cells. Data are means ± SD (n = 9, *N* = 3 for 5 BV; n = 6, *N* = 3 for + α_2_-δ and + γ1) and were analyzed by one-way ANOVA with Dunnett’s multiple comparisons test. “n” is the number of wells and “*N*” is the number of independent experiments.

Skeletal muscle Cav1.1 forms a complex with three auxiliary subunits, β1a, α_2_-δ and γ_1_ (Catterall, 2000). α_2_-δ and γ_1_ may play modulatory roles (Flucher et al., 2005). Additional expression of α_2_-δ had no effect on high [K^+^]-induced Ca^2+^ release (**Fig. 2 D, E**). Expression of γ1 did not significantly affect high [K^+^]-induced Ca^2+^ release at 20 mM or lower [K^+^] but it reduced the fluorescence change at higher [K^+^] concentrations (**Fig. 2 D, E**). In γ_1_ cells, R-CEPIA1er fluorescence decreased but increased again with time, suggesting inactivation of the Ca^2+^ release (**Fig. 2 A**). These findings are consistent with previous reports of the effects of α_2_-δ (Obermair et al., 2005) and γ_1_ (Ursu et al., 2004) on DICR. Therefore, our reconstituted platform can reproduce DICR in HEK293 cells and is valid for evaluating disease-causing mutations or drugs.

We also tested the importance of Kir2.1 on the reconstituted DICR platform. We measured membrane potential of cells under current-clamp conditions using pipette solution containing ∼145 mM [K^+^] (**Fig. 3 A**). The membrane potential of 5 BV-infected cells was -71 ± 3 mV in normal Krebs solution and became depolarized to -14 ± 3 mV by high [K^+^] solution containing 50 mM [K^+^]. In contrast, membrane potential was already depolarized in the cells without Kir2.1 (−18 ± 2 mV) in normal Krebs solution, which is consistent with the previous report(Kirkton and Bursac, 2011). Without Kir2.1, R-CEPIA1er fluorescence intensity was substantially reduced and high [K^+^]-induced Ca^2+^ release was completely abolished (**Fig. 3 B**). We quantified resting ER [Ca^2+^] by determining the maximum fluorescence intensity of R-CEPIA1er using ionomycin/Ca^2+^ (**Fig. 3 C**). The resting ER [Ca^2+^] was severely reduced in cells without Kir2.1 (**Fig. 3 D**). These results suggest that persistently activated DICR activity under the depolarized conditions causes ER Ca^2+^ depletion. In support of this, further removal of β1a or Stac3, which completely abolishes DICR activity, restored the ER [Ca^2+^] (**Fig. 3 D**). Therefore, Kir2.1 is indispensable for our reconstituted DICR platform.

**Figure 3.**
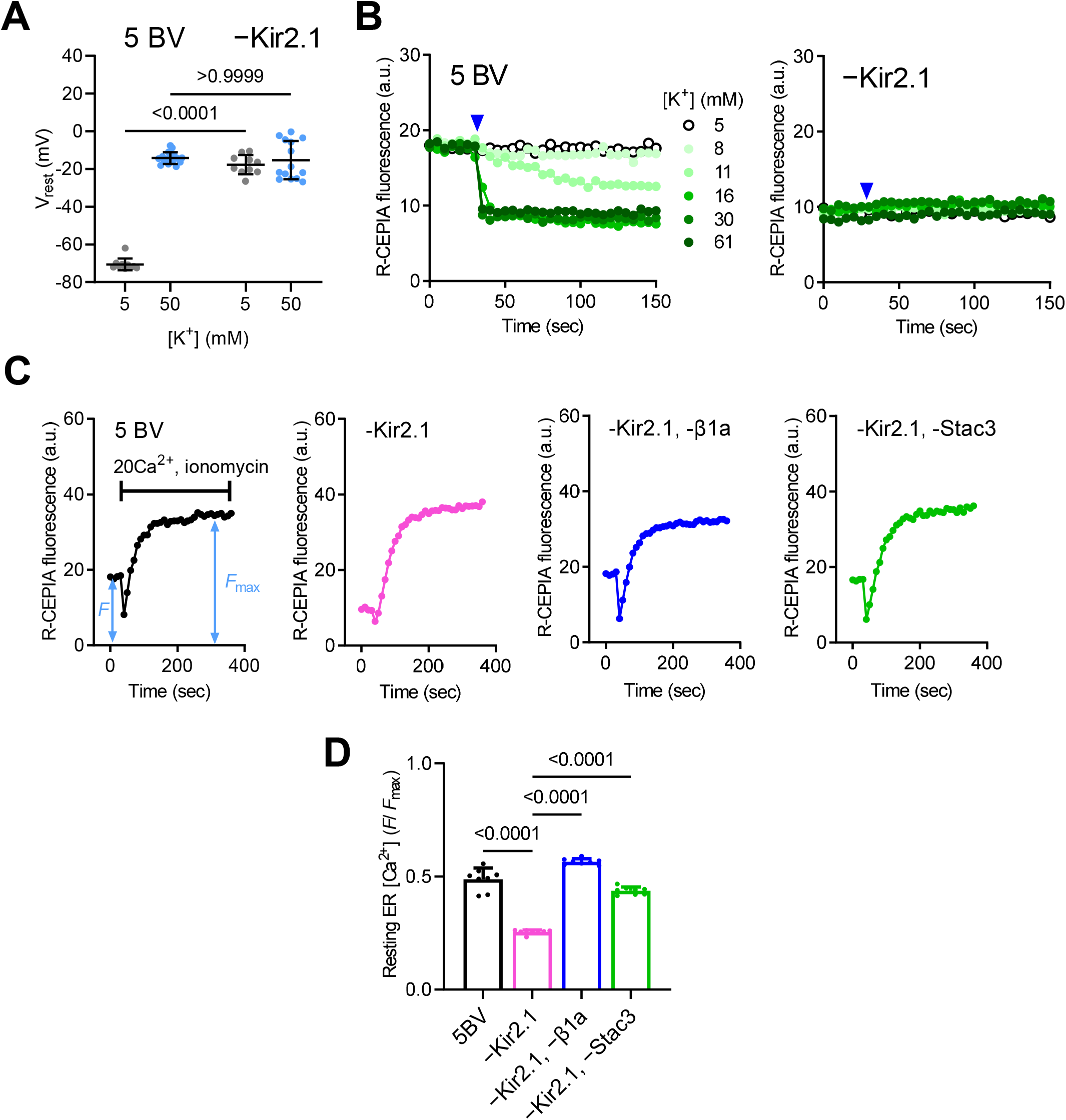
Importance of Kir2.1 on DICR platform. **(A)** Resting membrane potential of cells infected with 5 BV (left) or BV without Kir2.1 (−Kir2.1) (right). Data are means ± SD (n = 10, *N* = 3 for 5 mM [K^+^] with 5 BV; n = 16, *N* = 3 for 50 mM [K^+^] with 5 BV; n = 10, *N* = 3 for 5 mM [K^+^] with -Kir2.1 and n = 14, *N* = 3 for 50 mM [K^+^] with -Kir2.1) and were analyzed by two-way ANOVA with Bonferroni’s multiple comparisons test. **(B)** Typical results of R-CEPIA1er fluorescence in cells infected with 5 BV (left) and BV without Kir2.1 (right). High [K^+^] solution ranging from 5–61 mM (symbols shown in left) was applied at 30 sec after starting (blue arrowheads). **(C)** Typical results of ER [Ca^2+^] measurement in cells with 5 BV (left), without Kir2.1, without Kir2.1 and β1a, and without Kir2.1 and Stac3. **(D)** Resting ER [Ca^2+^] in cells infected with 5 BV, BV without Kir2.1 (−Kir2.1), Kir2.1 and β1a (−Kir2.1, -β1a), or Kir2.1 and Stac3 (−Kir2.1, -Stac3). Data are means ± SD (n = 8, *N* = 3) and were analyzed by one-way ANOVA with Dunnett’s multiple comparisons test. “n” is the number of cells (A) or wells (D) and “*N*” is the number of independent experiments.

### Evaluation of disease-causing mutations in Cav1.1

Using the reconstituted DICR platform, we evaluated mutations in Cav1.1 which are implicated in various muscle diseases (Beam et al., 2017; Flucher, 2020). We initially tested known autosomal dominant MH mutations, R174W (Carpenter et al., 2009), R1086H (Stewart et al., 2001), and T1354S (Pirone et al., 2010) (**Fig. 4 A**). Ca^2+^ non-conducting N617D mutation was also introduced in wild type (WT) and the mutant Cav1.1s. To observe the effect of mutations clearly, we tested them in homozygous state without WT. The expression of mutant Cav1.1s was confirmed by Western blot (**Fig. S1 C**). We found that ER [Ca^2+^] was severely reduced for R1086H (**Fig. 4 B**). In support of this, R1086H exhibited lower initial R-CEPIA1er fluorescence intensity and smaller fluorescence change in response to high [K^+^] than those in WT or the other mutants (**Fig. 4 C**). We plotted [K^+^] dependence of DICR using the F/F_0_ corrected by ER [Ca^2+^] (**Fig. 4 D**). This clearly indicates changes in ER [Ca^2+^] in each WT or mutant cell. The change in the corrected F/F_0_ was substantially smaller in R1086H (**Fig. 4 E**), being consistent with initial depletion of ER [Ca^2+^]. Interestingly, R1086H exhibited significantly lower EC_50_ value for [K^+^] (8.3 ± 0.3 mM) compared to WT (10.8 ± 0.7 mM), indicating an enhanced [K^+^]-dependence (**Fig. 4 F**). No or only minor changes in ER [Ca^2+^] and [K^+^] dependence were observed for R174W or T1354S cells (**Fig. 4 B–F**). We anticipated that the reduction in ER [Ca^2+^] in R1086H cells was caused by activation of Cav1.1 at resting membrane potential because of its hypersensitivity. To test the hypothesis, we hyperpolarized the membrane potential which reduced the activation of Cav1.1. Reduction of [K^+^] in the Krebs solution from 5 mM to 2.5 mM successfully hyperpolarized the membrane potential by ∼15 mV (**Fig. S4 A**). This substantially increased ER [Ca^2+^] in R1086H cells to a level comparable to that in WT cells (**Fig. S4 B**). R1086H cells exhibited a clear greater sensitivity to [K^+^] (5.9 ± 0.2 mM) than WT cells (8.3 ± 0.5 mM) (**Fig. S4 C–E**), supporting our hypothesis of R1086H hyperactivation.

**Figure 4.**
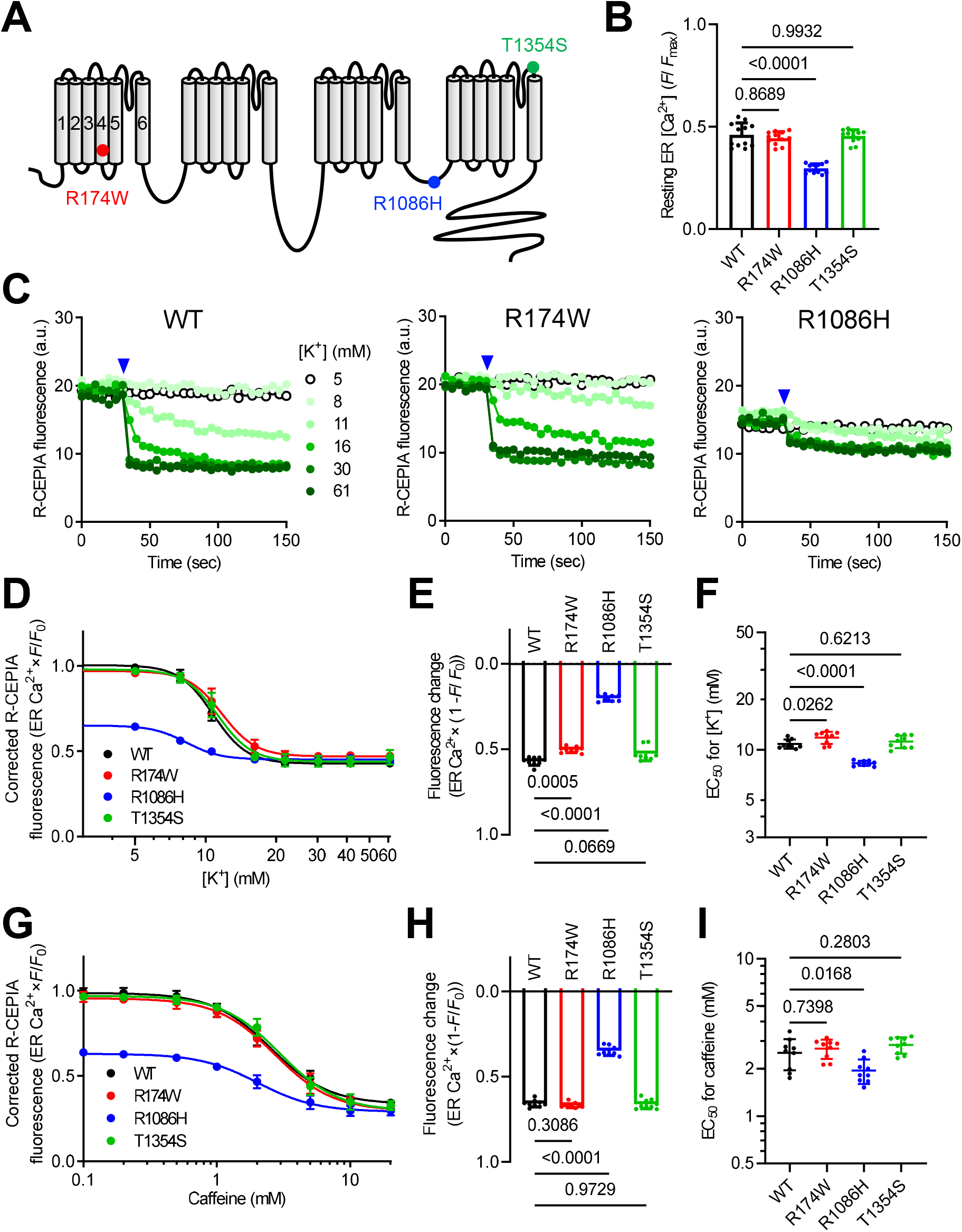
Effect of MH mutations in Cav1.1 on DICR activity. **(A)** Schematic drawing of the location of MH mutations (R174W, R1086H and T1354S) in Cav1.1. Note that WT and mutant Cav1.1s also carry N617D Ca^2+^ non-conducting mutation. **(B)** Resting ER [Ca^2+^] in cells expressing WT and mutant Cav1.1. Data are means ± SD (n = 12; *N* = 3) and were analyzed by one-way ANOVA with Dunnett’s multiple comparisons test. **(C)** Typical results of R-CEPIA1er fluorescence in cells infected with 5 BV carrying WT (left), R174W (center) and R1086H (right) Cav1.1. High [K^+^] solution ranging from 5–61 mM (symbols shown in left) was applied at 30 sec after starting (blue arrowheads). **(D)** [K^+^] dependence of R-CEPIA1er fluorescence corrected by ER [Ca^2+^] in WT (black), R174W (red), R1086H (blue) and T1354S (green) Cav1.1 cells. Data are means ± SD (n = 9, *N* = 3). **(E, F)** Fluorescence change by 61 mM [K^+^] corrected by ER Ca^2+^ (**E**) and EC_50_ values for [K^+^] (**F**). Data are means ± SD (n = 9, *N* = 3) and were analyzed by one-way ANOVA with Dunnett’s multiple comparisons test. **(G)** Caffeine dependence of R-CEPIA1er fluorescence corrected by ER Ca^2+^ in WT (black), R174W (red), R1086H (blue) and T1354S (green) Cav1.1 cells. Data are means ± SD (n = 9, *N* = 3). **(H, I)** Fluorescence change by 20 mM caffeine corrected by ER Ca^2+^ (**H**) and EC_50_ values for caffeine (**I**). Data are means ± SD (n = 9, *N* = 3) and were analyzed by one-way ANOVA with Dunnett’s multiple comparisons test. “n” is the number of wells and “*N*” is the number of independent experiments.

Previous functional characterization of these Cav1.1 mutants consistently showed enhanced caffeine sensitivity, i.e., acceleration of Ca^2+^-induced Ca^2+^ release (CICR) (Weiss et al., 2004; Pirone et al., 2010; Eltit et al., 2012). We tested caffeine dependence of these mutants in our platform. Caffeine released Ca^2+^ from WT cells in a dose-dependent manner (**Fig. 4 G**). R1086H cells exhibited a reduced fluorescence change by 20 mM caffeine (**Fig. 4 H**) and a significantly smaller EC_50_ (**Fig. 4 I**), indicating an enhanced caffeine sensitivity. Caffeine dependence was not changed in R174W or T1354S cells (**Fig. 4 G–I**).

We next examined the effect of myopathy-related mutations in Cav1.1 on DICR activity. We chose four mutations, E100K, F275L, P742Q, and L1367V (Schartner et al., 2017), none of which have been functionally characterized (**Fig. 5 A**). Ca^2+^ non-conducting N617D mutation was also introduced in the mutant Cav1.1s. and we tested these mutations in homozygous states without WT. The expression of mutant Cav1.1s was confirmed by Western blot (**Fig. S1 C**). ER [Ca^2+^] was not changed by any of these mutations (**Fig. 5 B**). DICR activity was completely lost in the F275L mutant and there was a substantial rightward shift in [K^+^] dependence for P742Q (**Fig. 5 C, D**). Whereas fluorescence change by 61 mM [K^+^] was not changed (**Fig. 5 E**), the EC_50_ value was threefold higher in P742Q (33.2 ± 1.0 mM) compared with WT (10.3 ± 0.9 mM) (**Fig. 5 F**). The other two mutations (E100K and L1367V) had no significant effects on [K^+^] dependence (**Fig. 5 D-F**). We also tested the effects of these mutations on caffeine-induced Ca^2+^ release. No substantial changes were observed with any of the mutations (**Fig. 5 G–I**).

**Figure 5.**
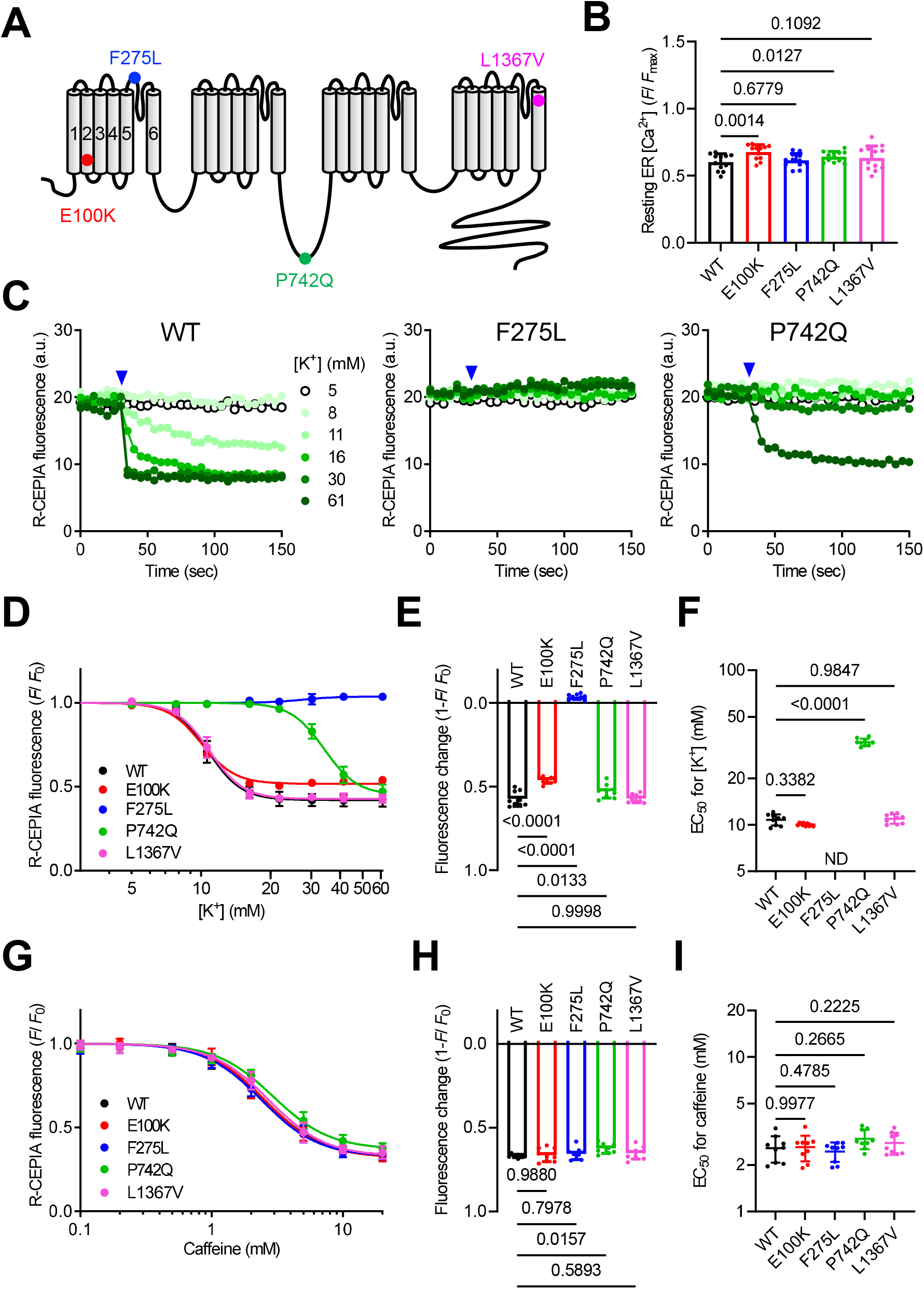
Effect of myopathy mutations in Cav1.1 on DICR activity. **(A)** Schematic drawing of the location of myopathy mutations (E100K, F275L, P742Q and L1367V) in Cav1.1. Note that WT and mutant Cav1.1s also carry N617D Ca^2+^ non-conducting mutation. **(B)** Resting ER [Ca^2+^] in cells expressing WT and mutant Cav1.1. Data are means ± SD (n = 12, *N* = 3) and were analyzed by one-way ANOVA with Dunnett’s multiple comparisons test. **(C)** Typical results of R-CEPIA1er fluorescence in cells infected with 5 BV carrying WT (left), F275L (center) and P742Q (right) Cav1.1. High [K^+^] solution ranging from 5–61 mM (symbols shown in left) was applied at 30 sec after starting (blue arrowheads). **(D)** [K^+^] dependence of R-CEPIA1er fluorescence (F/F_0_) in WT (black), E100K (red), F275L (blue), P742Q (green) and L1367V (magenta) Cav1.1 cells. Data are means ± SD (n = 9, *N* = 3). **(E, F)** Fluorescence change by 61 mM [K^+^] (**E**) and EC_50_ values for [K^+^] (**F**). Data are means ± SD (n = 9, *N* = 3) and were analyzed by one-way ANOVA with Dunnett’s multiple comparisons test. **(G)** Caffeine dependence of R-CEPIA1er fluorescence (F/F_0_) in WT (black), E100K (red), F275L (blue), P742Q (green) and L1367V (magenta) Cav1.1 cells. Data are means ± SD (n = 9, *N* = 3). **(H, I)** Fluorescence change by 20 mM caffeine (**H**) and EC_50_ values for caffeine (**I**). Data are means ± SD (n = 9, *N* = 3) and were analyzed by one-way ANOVA with Dunnett’s multiple comparisons test. “n” is the number of wells and “*N*” is the number of independent experiments.

### Effect of drugs on DICR activity

The reconstituted DICR platform is expected to be useful for screening drugs for muscle diseases. To test this possibility, we examined the effects of known DICR modulators. We initially tested three RyR1 inhibitors (**Fig. 6 A**). Dantrolene is a well-known RyR1 inhibitor that is clinically used for MH (Riazi et al., 2018). Cpd1 is a novel potent RyR1-selective inhibitor that we recently developed (Mori et al., 2019; Yamazawa et al., 2021). Procaine inhibits CICR but not DICR in frog skeletal muscle (Thorens and Endo, 1975; Kashiyama et al., 2010). Dantrolene (10 µM) and Cpd1 (3 µM) caused rightward shifts in [K^+^] dependence compared with the control (**Fig. 6 B, C**). The maximum response by 61 mM [K^+^] was not changed (**Fig. 6 D**), but EC_50_ value was significantly higher in dantrolene (13.1 ± 0.5 mM) or Cpd1 (13.5 ± 0.5 mM) compared to control (10.4 ± 0.3 mM) (**Fig. 6 E**). These findings indicate that dantrolene and Cpd1 inhibit DICR. This is consistent with the suppression of twitch and tetanic tension in isolated muscles and with reduced *in vivo* muscle weakness by dantrolene (Meyler et al., 1979; Leslie and Part, 1981) or Cpd1 (Yamazawa et al., 2021). Procaine (5 mM), in contrast, did not inhibit DICR but slightly reduced EC_50_ value (9.0 ± 0.6 mM) (**Fig. 6 B-E**). All three compounds significantly shifted the caffeine dependence rightward (**Fig. 6 F**). The maximum response by 20 mM caffeine was not changed (**Fig. 6 G**), but EC_50_ values were increased by the drugs (1.9 ± 0.2, 3.0 ± 0.2, 3.1 ± 0.1, and 3.3 ± 0.4 mM for control, dantrolene, Cpd1, and procaine, respectively) (**Fig. 6 H**). These results indicate that dantrolene and Cpd1 inhibit both DICR and CICR, whereas procaine selectively inhibits CICR.

**Figure 6.**
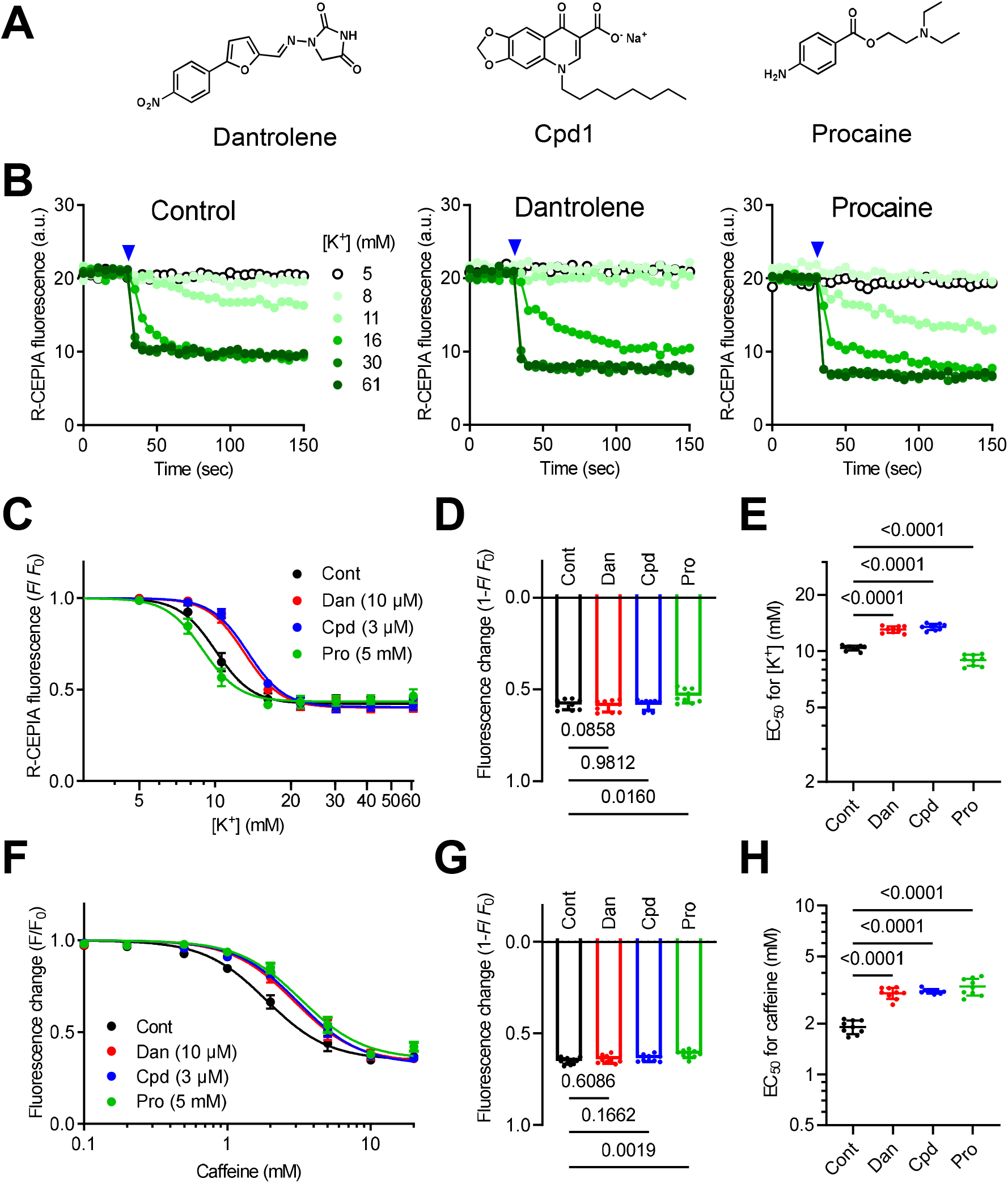
Effect of RyR1 inhibitors on DICR activity. **(A)** Chemical structures of dantrolene, Cpd1 and procaine. **(B)** Typical results of R-CEPIA1er fluorescence in cells without (Control, left) or with 10 μM dantrolene (center) or 5 mM procaine (right). High [K^+^] solution ranging from 5–61 mM (symbols shown in left) was applied at 30 sec after starting (blue arrowheads). **(C)** [K^+^] dependence of fluorescence change (F/F_0_) in cells without (Cont, black) or with 10 μM dantrolene (Dan, red), 3 μM Cpd1 (Cpd1, blue) or 5 mM procaine (Pro, green). Data are means ± SD (n = 9, *N* = 3). **(D, E)** Fluorescence change by 61 mM K^+^ (**D**) and EC_50_ values for [K^+^] (**E**). Data are means ± SD (n = 9, *N* = 3) and were analyzed by one-way ANOVA with Dunnett’s multiple comparisons test. **(F)** Caffeine dependence of R-CEPIA1er fluorescence (F/F_0_) in cells without (Control, black) or with 10 μM dantrolene (Dan, red), 3 μM Cpd1 (Cpd1, blue) or 5 mM procaine (Pro, green). Data are means ± SD (n = 9, *N* = 3). **(G, H)** Fluorescence change by 20 mM caffeine (**G**) and EC_50_ values for caffeine (**H**). Data are means ± SD (n = 9, *N* = 3) and were analyzed by one-way ANOVA with Dunnett’s multiple comparisons test. “n” is the number of wells and “*N*” is the number of independent experiments.

Lyotropic anions, such as perchlorate and thiocyanate, potentiate E-C coupling in skeletal muscle (Luttgau et al., 1983; Huang, 1986; Csernoch et al., 1987; Delay et al., 1990; Gonzalez and Rios, 1993). Perchlorate (10 mM) and thiocyanate (10 mM) significantly shifted the [K^+^] dependence leftward (**Fig. 7 A, B**). The EC_50_ values were significantly reduced (8.6 ± 0.3 and 9.2 ± 0.3 mM for perchlorate and thiocyanate, respectively) compared to control (10.6 ± 0.3 mM) without changing the maximum response (**Fig. 7 C, D**). In contrast, they did not affect caffeine dependence (**Fig. 7 E**); no significant changes were observed with the maximum response (**Fig. 7 F**) nor EC_50_ for caffeine (**Fig. 7 G**). These results indicate that lyotropic anions potentiate DICR but not CICR. This is consistent with previous findings showing that the potentiating effects of lyotropic anions are primarily caused by shifting voltage dependence of the charge movement toward more negative potentials (Luttgau et al., 1983; Huang, 1986; Csernoch et al., 1987; Delay et al., 1990; Gonzalez and Rios, 1993). Taken together, our reconstituted DICR platform reproduces the effects of known DICR modulators, indicating that it can be used to screen drugs that affect DICR.

**Figure 7.**
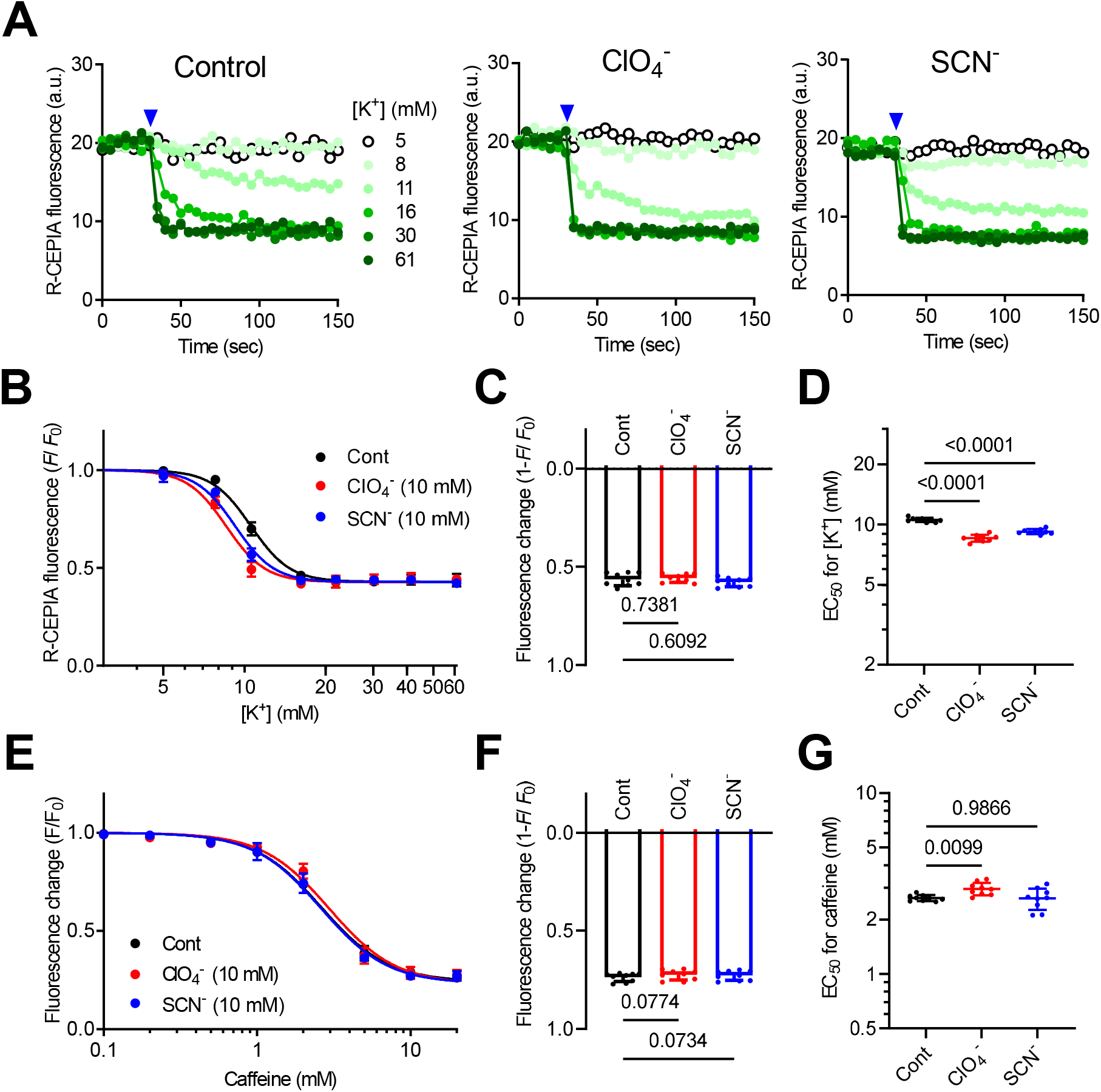
Effect of lyotropic anions on DICR activity. (**A**) Typical results of R-CEPIA1er fluorescence in cells in the absence (Control, left) and presence of 10 mM perchlorate (ClO ^-^, center) or 10 mM thiocyanate (SCN^-^, right). High [K^+^] solution ranging from 5–61 mM (symbols shown in left) was applied at 30 seconds after starting (blue arrowheads). **(B)** [K^+^] dependence of fluorescence change (F/F_0_) in cells in the absence (Cont, black) and presence of 10 mM perchlorate (ClO ^-^, red) or 10 mM thiocyanate (SCN^-^, blue). Data are means ± SD (n = 9, *N* = 3). **(C, D)** Fluorescence change by 61 mM K^+^ (**C**) and EC_50_ values for K ^+^ (**D**). Data are means ± SD (n = 9, *N* = 3) and were analyzed by one-way ANOVA with Dunnett’s multiple comparisons test. EC_50_ values were significantly reduced with lyotropic anions. **(E)** Caffeine dependence of R-CEPIA1er fluorescence (F/F_0_) in cells in the absence (Cont, black) and presence of 10 mM perchlorate (ClO ^-^, red) or 10 mM thiocyanate (SCN^-^, blue). Data are means ± SD (n = 9, *N* = 3). **(F, G)** Fluorescence change by 20 mM caffeine (**F**) and EC_50_ values for caffeine (**G**). Data are means ± SD (n = 9, *N* = 3) and were analyzed by one-way ANOVA with Dunnett’s multiple comparisons test. No significant changes were observed by lyotropic anions. “n” is the number of wells and “*N*” is the number of independent experiments.

## Discussion

In this study, we established a platform for DICR that was reconstituted in HEK293 cells expressing RyR1 and R-CEPIA1er. We made three key improvements to the original method by Perni et al. (Perni et al., 2017). First, the essential components were transduced using VSV-G pseudotyped baculovirus (BV), which can effectively infect a wide variety of mammalian cells without toxicity (Tani et al., 2001). This greatly increased the transduction efficiency; almost all cells expressed the essential components (**Fig. S1**). Second, depolarization of the plasma membrane was induced by high [K^+^] solution. This enabled us to simultaneously stimulate many cells (**Fig. 1, Fig. S2** and **movie S1**). For this purpose, we expressed Kir2.1, an inward-rectifying potassium channel, which effectively hyperpolarized the membrane potential (Kirkton and Bursac, 2011) (**Fig. 3**). Third, ER [Ca^2+^], instead of cytoplasmic [Ca^2+^], was measured to detect DICR. This is critically important to avoid contamination of signals from extracellular Ca^2+^ influx. Indeed, ER [Ca^2+^] was not changed by high [K^+^] depolarization in cells without essential components (**Fig. 1 A**), in contrast to substantial increases in cytoplasmic [Ca^2+^] (**Fig. S4 A**). These improvements allow quantitative measurements of DICR using a microplate reader.

In our reconstitution, four components, RyR1, Cav1.1, β1a, and Stac3, are essential for DICR; removal of each component totally abolished the DICR activity (**Fig. 2 A, B**). This is consistent with previous findings from mice lacking each component (Tanabe et al., 1988; Takeshima et al., 1994; Gregg et al., 1996; Nelson et al., 2013). In addition, the effects of α_2_-δ and γ_1_ auxiliary subunits on DICR activity corresponded with findings of previous reports (Ursu et al., 2004; Obermair et al., 2005) (**Fig. 2 C, D**). Therefore, our reconstituted platform successfully reproduces DICR in HEK293 cells. We found that a substantial DICR still occurred without JP2 (**Fig. 2 A, B**). This is in contrast to reports by Perni et al. (Perni et al., 2017; Perni and Beam, 2022), in which no voltage-gated Ca^2+^ release was observed in reconstituted cells without junctophilins. A possible reason for this is differences in expression methods. Our VSV-G BV is more effective at transduction than standard lipofection; therefore, sufficient proteins were expressed that might spontaneously interact with each other without junctophilins. Thus, our data suggest that junctophilins are not essential for DICR but may support formation of T-tubule-SR junction (Takeshima et al., 2000). It would be of interest whether the tetrads (structural signs for functional coupling between DHPR and RyR1) (Franzini-Armstrong, 2018) are formed in the cells expressing essential components without junctophilins.

We quantitatively evaluated the effect of disease-causing mutations or drugs using [K^+^] dependence. The EC_50_ value for [K^+^] in ER [Ca^2+^] measurement was ∼17 mM by calculation from the rapid fluorescence change (5 sec after stimulation) (**Fig. 1 C**). This is consistent with the value obtained from cytoplasmic [Ca^2+^] measurement (∼16 mM, **Fig. S3**) or those obtained from mouse skeletal muscle myotubes (∼21 mM) in high [K^+^]-induced Ca^2+^ transients (Yang et al., 2006; Lopez et al., 2018). However, since variations were large under the condition, we instead adopted the averaged fluorescence intensity for 125–150 sec after stimulation, which was more stable and accurate (**Fig. 1 C**). Although the EC_50_ value was reduced to ∼10 mM, the results reproduced the effects of disease-causing mutations (**Fig. 3–5**) or drugs (**Fig. 6, 7**), strongly indicating that the EC_50_ value for [K^+^] is a valid parameter for quantitative evaluation of DICR.

Mutations in Cav1.1 are implicated in various muscle diseases, including MH and myopathy (Beam et al., 2017; Flucher, 2020). Among three MH mutations (R174W, R1086H and T1354S), we demonstrated that R1086H, but not R174W or T1354S, significantly shifted the voltage dependence of DICR to more negative potentials (**Fig. 4 C–F**). This is consistent with previous findings using Cav1.1-deficient (dysgenic) myotubes (Weiss et al., 2004; Pirone et al., 2010; Eltit et al., 2012). All three mutations are reported to an increased sensitivity to caffeine-induced Ca^2+^ release compared with WT, indicating enhanced CICR activity (Weiss et al., 2004; Pirone et al., 2010; Eltit et al., 2012). However, we showed that caffeine sensitivity was increased only in R1086H (**Fig. 4 G–I**). A possible reason for this difference is the cells used. We used stable RyR1-overexpressing cells. This may cause an excess amount of ‘uncoupled’ RyR1, which might mask the enhanced sensitivity of caffeine dependence of the ‘coupled’ RyR1 by Cav1.1 mutations.

Most myopathy-related mutations identified in Cav1.1 have not been functionally characterized. We evaluated four mutations (E100K, F275L, P742Q and L1367V) found in patients with congenital myopathy (Schartner et al., 2017) (**Fig. 5**). To observe the effect of mutations clearly, we tested the mutations in homozygous states. We demonstrated that P742Q substantially inhibited DICR with a large rightward shifted [K^+^] dependence (**Fig. 5 C-F**). This proline reside in the II-III loop interacts with Stac3 (Wong King Yuen et al., 2017). Substitution of the proline to threonine, the residue found in the corresponding position of Cav1.2, diminished DICR in dysgenic myotubes (Kugler et al., 2004). Thus, the reduced voltage dependence of DICR might partly be involved in pathophysiology of the patients. E100K and L1367V had no or only minor impacts on the DICR activity (**Fig. 5 D-F**). However, our platform cannot detect rapid kinetics of the channel. Measurements of ionic currents and charge movements by patch-clamp method would further characterize these mutants. F275L is recessive because it was found in two compound heterozygous sisters carrying F275L and the nonsense mutation, p.L791Cfs*37 (Schartner et al., 2017). F275L completely abolished DICR without any changes in ER [Ca^2+^] or caffeine sensitivity (**Fig. 5 C-E**). F275L was expressed at a size similar to WT (**Fig. 1 C**) and ER [Ca^2+^] was preserved (**Fig. 5 B**). Thus, loss of DICR activity may not be due to loss of protein expression nor ER [Ca^2+^] depletion. Loss of DICR in F275L does not fit with a recessive mutation; if this is the case, patients will have no DICR activity. Further investigations are required for this mutation.

There are no specific treatments for most DICR-related diseases; therefore, there is an urgent need to develop novel drugs for these diseases (Agrawal et al., 2018; Flucher, 2020; Lawal et al., 2020). We here evaluated DICR modulators. Among three known RyR1 inhibitors, dantrolene and Cpd1 suppressed DICR (**Fig. 6**). This is consistent with the suppression of twitch and tetanic tensions of isolated muscles and of reduced *in vivo* muscle weakness by dantrolene (Meyler et al., 1979; Leslie and Part, 1981) and Cpd1 (Yamazawa et al., 2021). Interestingly, procaine did not suppress DICR (**Fig. 6**). Procaine inhibits CICR but not DICR in frog skeletal muscle (Thorens and Endo, 1975) and reconstituted frog α-RyR and β-RyR in RyR1-deficient myotubes (Kashiyama et al., 2010). Our data show similar procaine action in a mammalian system. We also demonstrated that lyotropic anions (perchlorate and thiocyanate) shifted the voltage-dependence of DICR to more negative potentials (**Fig. 7**). This reproduces previous findings using skeletal muscle fibers (Luttgau et al., 1983; Huang, 1986; Csernoch et al., 1987; Delay et al., 1990; Gonzalez and Rios, 1993). Interestingly, lyotropic anions did not affect caffeine dependence, suggesting no effect on CICR (**Fig. 7**). Therefore, our reconstituted DICR platform will also be useful for drug testing and screening for novel drugs that affect DICR.

## Limitations and future perspective

The present reconstituted DICR platform has several limitations to be considered. First, we overexpressed the essential components into HEK293 cells. Although this achieved efficient formation of the DICR machinery, proteins were not only localized to plasma membrane but also within cells except for in nuclei (Fig. S1 A). This is far from skeletal muscle triads. The excess amount of uncoupled proteins might affect the results. Peripheral localization of the components, e.g., RyR1 and Cav1.1, has been shown in the reconstituted cells by Perni et al. (Perni et al., 2017; Perni and Beam, 2022). In our platform, protein expression level can be easily controlled by amount of BV. Finding of the best amount and combination ratio of each component would improve the situation.

Second, we measured DICR using long-lasting (∼150 sec) high K^+^ depolarization. This provided efficient stimulation of the reconstituted cells but is markedly different from the rapid Ca^2+^ release kinetics (∼few ms) that occur during an action potential in muscle. Parallel experiments using patch-clamp method is necessary to characterize the mutants. Our reconstituted cells are applicable to single cell Ca^2+^ measurements combined with patch-clamp method. This would measure the rapid kinetics of Ca^2+^ release by direct membrane depolarization.

Third, we expressed the minimum essential components (RyR1, Cav1.1, β1a, Stac3, JP2 and Kir2.1) for functional reconstitution of DICR. This is simple but different from the skeletal muscle environment, in which a number of proteins (e.g., triadin, junctin, calsequestrin, FKBP12, JP45 etc) associates and regulates the DICR machinery. Drug screening with reconstitution by the minimum essential components might miss the compounds that act on the associated proteins. Since VSV-G BV system easily expresses additional proteins (e.g., α_2_-δ or γ_1_ subunit, see Fig. 2 D), it is possible to express the associated proteins alone or in combination in our DICR platform. It would also be interesting to express voltage-gated sodium and potassium channels to support electrically-evoked Ca^2+^ release near future.

## Conclusions

In summary, we established a reconstituted DICR platform in HEK293 cells. The platform is highly efficient and quantitative thus will be useful for both evaluation of disease-causing mutations and the development of novel drugs for DICR -related diseases. Since the procedure is simple and reproducible, the reconstituted DICR platform will be a powerful tool for accelerating diagnosis and treatment of these diseases.

## Acknowledgments

We thank Ikue Hiraga and the Laboratory of Proteomics and Biomolecular Science, Biomedical Research Core Facilities, Juntendo University Graduate School of Medicine, for technical assistance. We thank Jeremy Allen, PhD, from Edanz (https://jp.edanz.com/ac) for editing a draft of this manuscript. This work was supported by JSPS KAKENHI (19H03404 and 22H02805 to T.M., 19K07105 and 22K06652 to N.K., 20K11368 to T.K.), the Platform Project for Supporting Drug Discovery and Life Science Research (Basis for Supporting Innovative Drug Discovery and Life Science Research (BINDS)) from the Japan Agency for Medical Research and Development (AMED) (JP20am0101080 to T.M. and N.K. and JP20am0101098 to H.K.), an Intramural Research Grant for Neurological and Psychiatric Disorders from the National Center of Neurology and Psychiatry (2-5 to T.M.), the Vehicle Racing Commemorative Foundation (6237 and 6303 to T.M.), and the Cooperative Research Project of Research Center for Biomedical Engineering (to H.K.). The authors declare no competing financial interests.

## Author contributions

T. Murayama, N. Kurebayashi, S. Okazaki, K. Yamashiro carried out the cell biological experiments; T. Numaga-Tomita and M. Yamada performed patch-clamp experiments; T. Nakada, S. Mori, R. Ishida, and H. Kagechika provided experimental tools; T. Murayama, N. Kurebayashi, T. Numaga-Tomita, T. Kobayashi, T. Nakada, M. Yamada, and T. Sakurai analyzed the experimental data; T. Murayama, N. Kurebayashi, T. Numaga-Tomita and M. Yamada wrote the manuscript. All authors reviewed and approved the manuscript.

## Figure legends

Figure S1 **Expression of essential components for DICR machinery using VSV-G pseudotyped BV. (A)** Immunofluorescent detection of Cav1.1, β1a, Stac3, JP2 and Kir2.1 in 5 BV (left) and No BV (right) cells. Cells were labeled with antibodies to each component, followed by Alexa594-labeled anti-mouse IgG. Scale bar, 20 μm. Note that red fluorescence was specifically observed with 5 BV cells. **(B)** Western blots of essential components. Lysates from No BV and 5 BV cells were separated by SDS-PAGE and probed with antibodies to each component. Calnexin (CLNX) was used as a loading control. (**C**) Western blot of Cav1.1 mutants. Lysates from 5 BV cells carrying WT, R174W, R1086H, T1354S, E100K, F275L, P742Q and L1367V Cav1.1 were separated by SDS-PAGE and probed with anti-Cav1.1 antibody. Calnexin (CLNX) was used as loading control.

Figure S2 **Visualization of DICR by laser scanning confocal microscopy. (A)** Typical results of time-lapse R-CEPIA1er fluorescence measurement of individual 5 BV-infected RyR1/R-CEPIA1er HEK293 cells using a laser scanning confocal microscope (see also Movie S1). Cells were incubated with normal Krebs solution and high [K^+^] (50 mM) solution was perfused during measurements (arrow). Note that R-CEPIA1er fluorescence was transiently decreased by high [K^+^] solution and recovered after replacement with normal Krebs solution. **(B)** Percent responding cells to high [K^+^] (50 mM) or caffeine (10 mM) solution. 100 cells were randomly picked from each of No BV or 5 BV cells. Whereas 95% of 5 BV cells responded to high [K^+^], less than 1% of No BV cells responded. All the cells responded to caffeine. Data are means ± SD (n = 3, *N* = 3). “n” is the number of cells and “*N*” is the number of independent experiments.

Figure S3 **Measurement of DICR with cytoplasmic [Ca^2+^]. (A)** Typical results of time-lapse G-GECO1.1 fluorescence measurement in No BV (left) and 5 BV (right) cells. High [K^+^] solution ranging from 5–61 mM (shown at right) was applied at 30 sec after starting (blue arrows). Note that a substantial Ca^2+^ transient was observed in No BV, indicating the existence of endogenous Ca^2+^ influx pathways by depolarization. **(B)** [K^+^] dependence of fluorescence change (F/F_0_) in No BV (black) or 5 BV cells (red). F/F_0_ was obtained by normalizing the fluorescence signal at 35 sec (F, grey lines in **A**) to the averaged signals for the first 25 sec (F_0_). Data are means±SD (n = 6, *N* = 3). “n” is the number of wells and “*N*” is the number of independent experiments.

Figure S4 **Effect of the Cav1.1 R1086H mutation on DICR activity at 2.5 mM [K**^+^**]. (A)** Resting membrane potential of 5 BV cells in 5 mM or 2.5 mM [K^+^] Krebs solution. Data are the median with individual data points (n = 10, *N* = 3 for 5 mM [K^+^] and n = 20, *N* = 3 for 2.5 mM [K^+^]) and were analyzed by the unpaired two-tailed t test with the Mann-Whitney test. **(B)** Resting ER [Ca^2+^] in WT (black) and R1086H (blue) Cav1.1 cells. Data are means ± SD (n = 12, *N* = 3) and were analyzed by the two-tailed T-test with the Mann Whitney test. **(C)** [K^+^] dependence of R-CEPIA1er fluorescence (F/F_0_) in WT (black) and R1086H (blue) Cav1.1 cells. Data are means ± SD (n = 9, *N* = 3). **(D, E)** Fluorescence change by 61 mM [K^+^] (**D**) and EC_50_ values for [K^+^] (**E**). Data are means ± SD (n = 9, *N* = 3) and were analyzed by the two-tailed T-test with the Mann Whitney test. “n” is the number of cells (A) or wells (B-E) and “*N*” is the number of independent experiments.

