## Supplementary figures for "Reconstituted depolarization-induced Ca^2+^ release platform for skeletal muscle disease mutation validation and drug discovery"

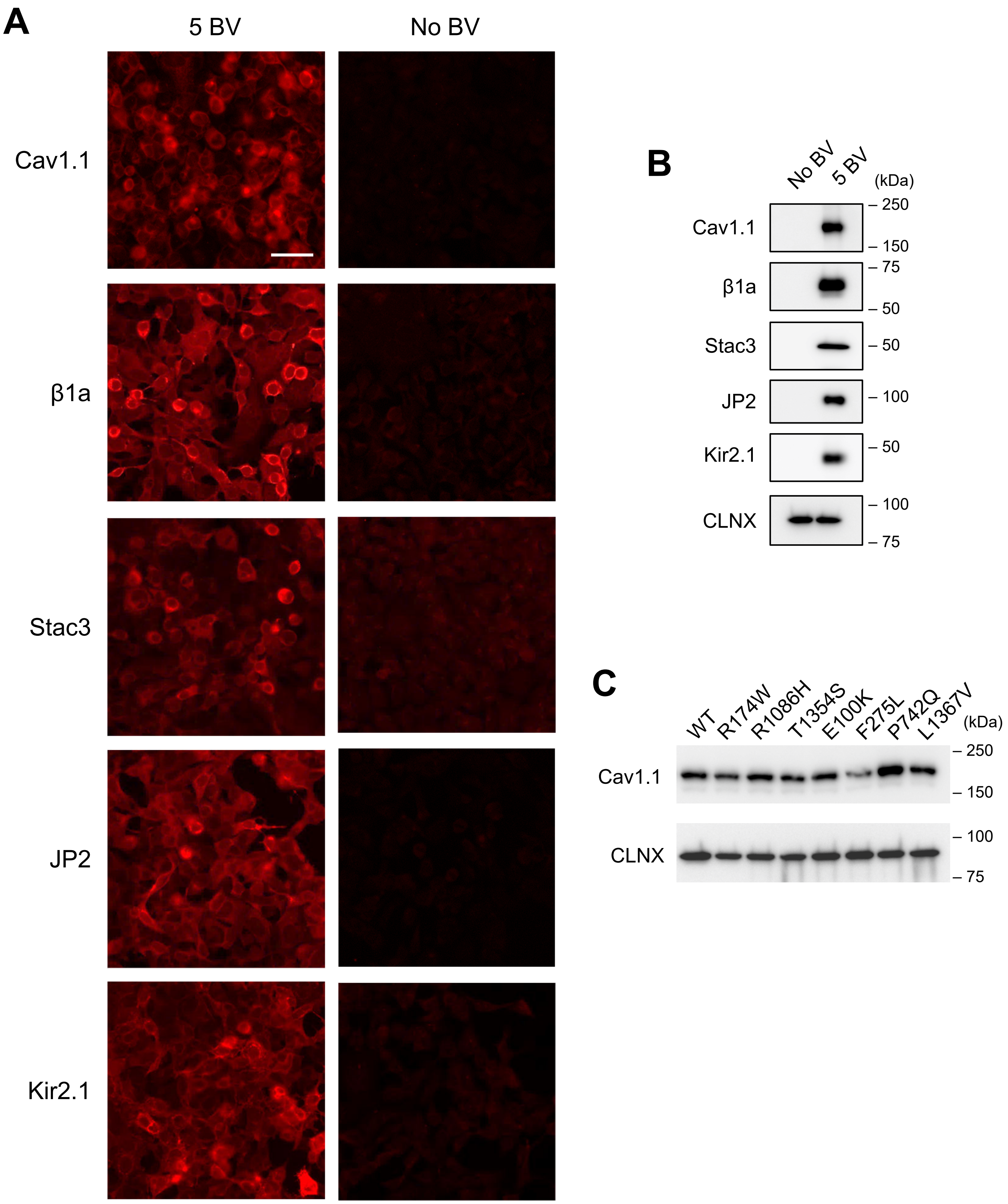


Figure S1 **Expression of essential components for DICR machinery using VSV-G pseudotyped BV.** **(A)** Immunofluorescent detection of Cav1.1, β1a, Stac3, JP2 and Kir2.1 in 5 BV (left) and No BV (right) cells. Cells were labeled with antibodies to each component, followed by Alexa594-labeled anti-mouse IgG. Scale bar, 20 μm. Note that red fluorescence was specifically observed with 5 BV cells. **(B)** Western blots of essential components. Lysates from No BV and 5 BV cells were separated by SDS-PAGE and probed with antibodies to each component. Calnexin (CLNX) was used as a loading control. (**C**) Western blot of Cav1.1 mutants. Lysates from 5 BV cells carrying WT, R174W, R1086H, T1354S, E100K, F275L, P742Q and L1367V Cav1.1 were separated by SDS-PAGE and probed with anti-Cav1.1 antibody. Calnexin (CLNX) was used as loading control.

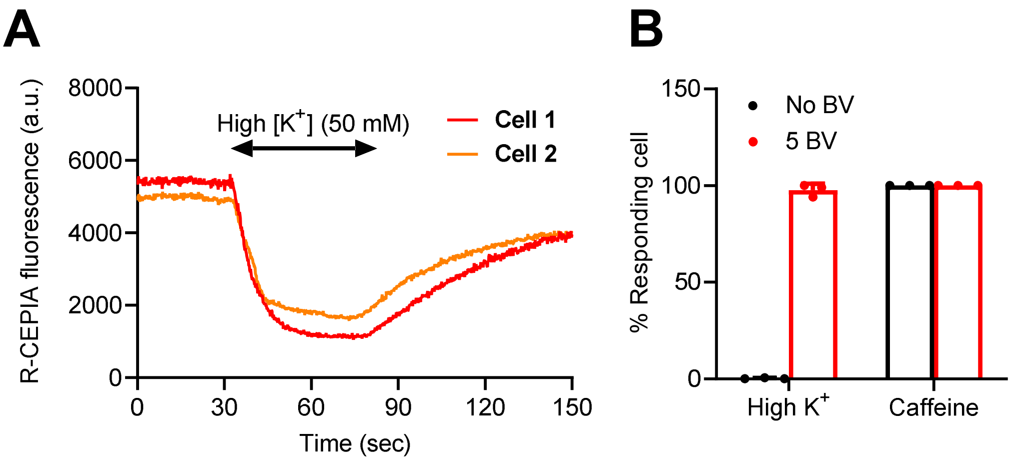


Figure S2 **Visualization of DICR by laser scanning confocal microscopy. (A)** Typical results of time-lapse R-CEPIA1er fluorescence measurement of individual 5 BV-infected RyR1/R-CEPIA1er HEK293 cells using a laser scanning confocal microscope (see also Movie S1). Cells were incubated with normal Krebs solution and high [$K^+$] (50 mM) solution was perfused during measurements (arrow). Note that R-CEPIA1er fluorescence was transiently decreased by high [$K^+$] solution and recovered after replacement with normal Krebs solution. **(B)** Percent responding cells to high [$K^+$] (50 mM) or caffeine (10 mM) solution. 100 cells were randomly picked from each of No BV or 5 BV cells. Whereas 95% of 5 BV cells responded to high [$K^+$], less than 1% of No BV cells responded. All the cells responded to caffeine. Data are means ± SD (n = 3, *N* = 3). "n" is the number of cells and "*N*" is the number of independent experiments.

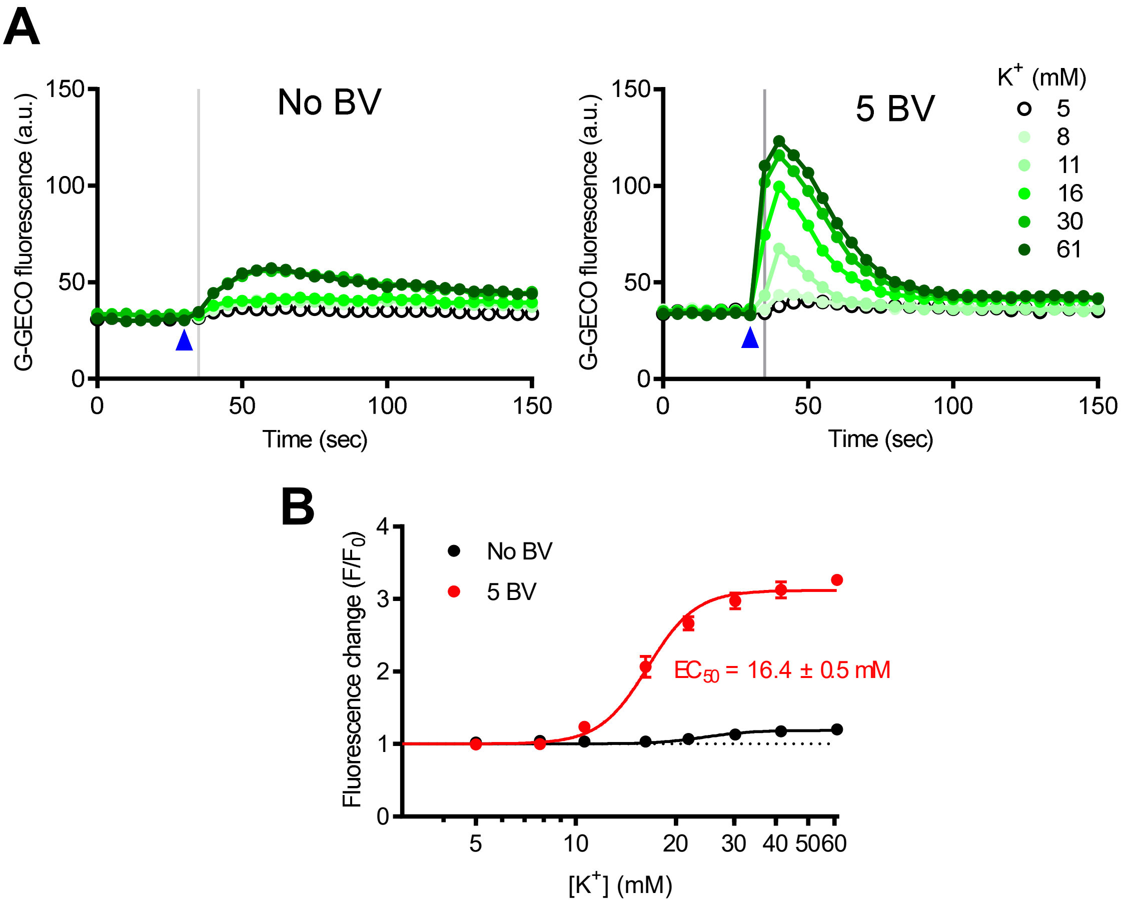


Figure S3 **Measurement of DICR with cytoplasmic [$Ca^{2+}$]. (A)** Typical results of time-lapse G-GECO1.1 fluorescence measurement in No BV (left) and 5 BV (right) cells. High [$K^+$] solution ranging from 5–61 mM (shown at right) was applied at 30 sec after starting (blue arrows). Note that a substantial $Ca^{2+}$ transient was observed in No BV, indicating the existence of endogenous $Ca^{2+}$ influx pathways by depolarization. **(B)** [$K^+$] dependence of fluorescence change ($F/F_0$) in No BV (black) or 5 BV cells (red). $F/F_0$ was obtained by normalizing the fluorescence signal at 35 sec (F, grey lines in **A**) to the averaged signals for the first 25 sec ($F_0$). Data are means±SD (n = 6, *N* = 3). "n" is the number of wells and "*N*" is the number of independent experiments.

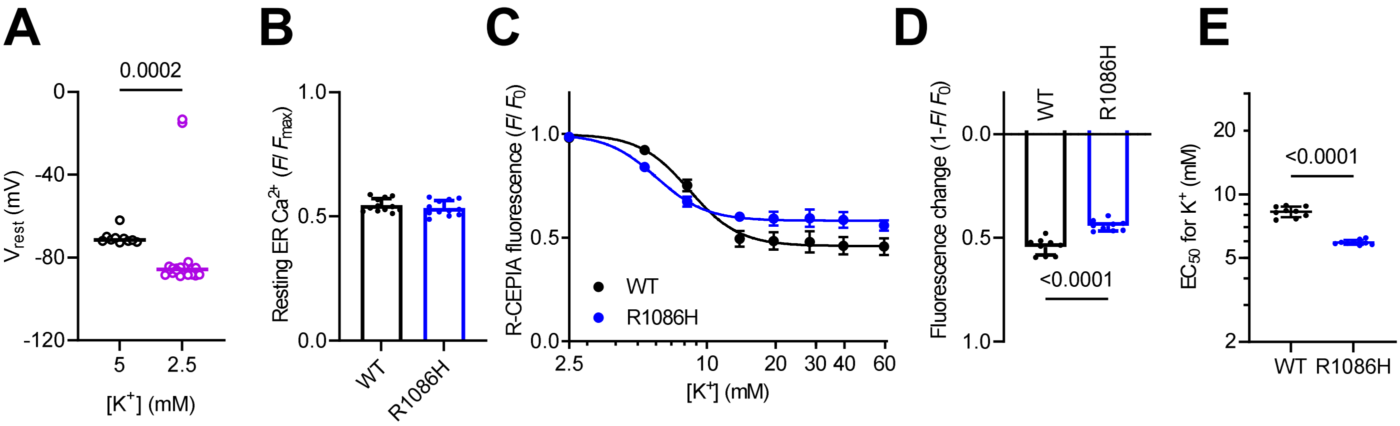


Figure S4 **Effect of the Cav1.1 R1086H mutation on DICR activity at 2.5 mM [K$^+$]. (A)** Resting membrane potential of 5 BV cells in 5 mM or 2.5 mM [K$^+$] Krebs solution. Data are the median with individual data points (n = 10, *N* = 3 for 5 mM [K$^+$] and n = 20, *N* = 3 for 2.5 mM [K$^+$]) and were analyzed by the unpaired two-tailed t test with the Mann-Whitney test. **(B)** Resting ER [Ca$^{2+}$] in WT (black) and R1086H (blue) Cav1.1 cells. Data are means ± SD (n = 12, *N* = 3) and were analyzed by the two-tailed T-test with the Mann Whitney test. **(C)** [K$^+$] dependence of R-CEPIA1er fluorescence (F/F$_0$) in WT (black) and R1086H (blue) Cav1.1 cells. Data are means ± SD (n = 9, *N* = 3). **(D, E)** Fluorescence change by 61 mM [K$^+$] (**D**) and EC$_{50}$ values for [K$^+$] (**E**). Data are means ± SD (n = 9, *N* = 3) and were analyzed by the two-tailed T-test with the Mann Whitney test. "n" is the number of cells (A) or wells (B-E) and "*N*" is the number of independent experiments.
